# A single- and paired-pulse TMS-EEG investigation of the N100 and long interval cortical inhibition in autism spectrum disorder

**DOI:** 10.1101/2021.01.06.425653

**Authors:** Melissa Kirkovski, Aron T. Hill, Nigel C Rogasch, Takashi Saeki, Bernadette M. Fitzgibbon, Joel Yang, Michael Do, Peter H Donaldson, Natalia Albein-Urios, Paul B Fitzgerald, Peter G Enticott

**Affiliations:** Cognitive Neuroscience Unit, School of Psychology, Deakin University, Geelong, Victoria, Australia; Brain, Mind and Society Research Hub, School of Psychological Sciences and Turner Institute for Brain and Mental Health, Monash University, Melbourne, Australia; Monash Biomedical Imaging, Monash University, Melbourne, Australia; South Australian Health and Medical Research Institute (SAHMRI), Adelaide, Australia; Discipline of Psychiatry, School of Medicine, University of Adelaide, Adelaide, Australia; Department of Psychiatry, Yokohama City University School of Medicine, Yokohama, Japan; Epworth Centre for Innovation in Mental Health, Epworth HealthCare and Central Clinical School, Monash University, Melbourne, Australia; Respiratory and Sleep Medicine, The Royal Children’s Hospital, Melbourne, Australia

**Author notes:** **Correspondence:** Dr Melissa Kirkovski.

**Keywords:** autism, transcranial magnetic stimulation, electroencephalography, gamma-aminobutyric acid, N100, long interval cortical inhibition

## Abstract

**Background:** There is evidence to suggest a disruption of gamma-aminobutyric acid (GABA) in autism spectrum disorder (ASD), but findings are mixed. Concurrent electroencephalography and transcranial magnetic stimulation (TMS-EEG) provides a novel method by which to probe GABA-mediated cortical inhibition.

**Methods:** With a particular focus on GABAB-ergic mechanisms, we investigated the N100 peak of the TMS evoked potential (TEP), as well as long interval cortical inhibition (LICI_EEG_) in adults with ASD (n = 23; 12 female) without intellectual disability, and a neurotypical comparison group (n =22; 12 female) matched for age, sex, and IQ. Seventy-five single-(spTMS) and 75 paired-(ppTMS; 100 ms inter-stimulus-interval) pulses were applied to the right primary motor cortex (M1), right temporoparietal junction (TPJ), and right dorsolateral prefrontal cortex (DLPFC) while EEG was recorded from 20 scalp electrodes. Additionally, electromyography (EMG) was used to investigate corticospinal inhibition following ppTMS to M1 (LICI_EMG_).

**Results:** There were no group differences in the N100 amplitude or latency following spTMS. LICI outcomes following ppTMS, as measured by either EEG or EMG, also did not differ between groups. These findings were further supported by Bayesian analyses, which provided weak-moderate support for the null hypothesis.

**Limitations:** Data presented here reflect adults without intellectual disability, and the generalisability of these results is therefore limited.

**Conclusions:** The findings of this study argue against GABAB-ergic impairment in adults with ASD without intellectual disability, at least at the cortical regions examined. Further research investigating these mechanisms in ASD at various ages, with varying degrees of symptomatology, and at different brain sites is an important factor in understating the role of GABA in the neuropathophysiology of ASD.

## Background

Autism spectrum disorder (ASD) is a neurodevelopmental condition characterised by social impairment and the presence of restricted and repetitive patterns of behaviours and interests (1). An imbalance between cortical excitation and inhibition is often considered to underlie, at least in part, the neuropathophysiology of ASD (2, 3). Dysregulation of gamma-aminobutyric acid (GABA), the brain’s primary inhibitory neurotransmitter, is a key aspect of this theory (2–5). In line with this, neuroimaging studies using magnetic resonance spectroscopy (MRS) commonly indicate that GABA concentration is reduced among individuals with ASD compared to neurotypical controls. This reduction has been identified at various frontal/motor (6–11), temporal (8, 12, 13), occipital (14), and cerebellar (10) voxels. There are, however, also MRS reports of no differences in GABA indices between individuals with and without ASD at these, and other, sites (11, 15–18), or when using alternative approaches to quantifying GABA concentration (13).

Unlike MRS, which indexes metabolite concentration, transcranial magnetic stimulation (TMS) is a non-invasive brain stimulation (NIBS) technique capable of probing synaptic reactivity, and variations in TMS protocols can target and modulate different intra-cortical neural mechanisms. When applied to the motor cortex, with outcomes measured via electromyography (EMG), paired-pulse TMS (ppTMS) protocols whereby two TMS pulses are delivered in quick succession, can provide putative measures of GABA-ergic function, depending on the specific stimulation parameters used (19). Short interval cortical inhibition (SICI) paradigms, in which a sub-threshold conditioning pulse is followed (within 5 ms) by a supra-threshold test pulse (19, 20), is likely to result from activity at GABAA receptors (21, 22). Alternatively, and of particular relevance to the present study, long interval cortical inhibition (LICI), defined by an inter-stimulus interval (ISI) of 50-200 ms between two suprathreshold pulses (23), is associated with activity at GABAB receptors (24, 25).

This capacity to differentiate between GABAA- and GABAB-ergic receptor activity via TMS is of great potential therapeutic benefit in ASD. Based on the general understanding of reduced GABA in ASD, GABA agonists (e.g., bumetanide and baclofen, a selective GABAB-agonist) have been trialled as pharmacological interventions for the core symptoms of ASD (26–30). Knowledge derived from TMS, therefore, could have important implications for drug selection, and also for understanding the effects of such drugs at a neurophysiological level. One caveat of the technique, however, is that TMS has long been constrained to the primary motor cortex (M1) due to reliance on a peripheral EMG response as an indirect marker of corticospinal excitability.

Consequently, the findings of studies using TMS-EMG protocols are less directly generalisable to brain regions more frequently associated with the core symptoms of ASD. Recent advances in amplifier technology and signal analysis techniques have enabled the recording of electrophysiological activity, using electroencephalography (EEG), concurrent to TMS application (see 31, 32). This technique, termed TMS-EEG, allows for the investigation of non-motor cortical regions, including those more directly implicated in the neuropathophysiology of ASD. It must be noted, however, that there is evidence to suggest that SICI and LICI ppTMS protocols produce similar EEG outcomes, and which advises against using ppTMS-EMG effects to infer about ppTMS-EEG (33). Therefore, in this paper, we will differentiate between EMG and EEG based LICI outcomes (LICI_EMG_ and LICI_EEG_, respectively).

The N100 of the TMS-evoked potential (TEP) is a large negative deflection in the EEG trace typically observed ~90 – 140 ms post-TMS pulse. Given the timing of this component, it is likely that the N100 represents GABAB-ergic mechanisms (31). Further, long interval ppTMS protocols, which as mentioned above are GABAB receptor dependant (19, 24, 25), have also been shown to suppress later components (including the N100), but not earlier TEPs (34), which are typically associated with GABA_A_ receptor activity (35). MRS research also supports a role of GABA-ergic mechanisms in the generation of this component (36). Du and colleagues (36) report an association between the glutamate/GABA ratio derived from the medial prefrontal cortex and the N100 response recorded from the corresponding Fz EEG electrode. Specifically, increased GABA and decreased glutamate was associated with a smaller N100 component, while lower GABA and higher glutamate predicted larger N100 amplitudes. This relationship was not replicated between MRS derived glutamate/GABA indices from the medial prefrontal cortex and the N100 recorded from the motor cortex or the vertex (36). Finally, there is also pharmacological evidence to support the notion of GABAB receptor involvement in the N100 (35).

In ASD, research using TMS-EMG protocols to investigate GABA-ergic responses is limited, and the outcomes are variable. These studies have been reviewed in detail previously (3, 37). Briefly, there is some evidence of SICI_EMG_ deficits in specific ASD sub-types, rather than ASD broadly, based on earlier diagnostic differentiation between autistic and Asperger’s disorders (38, 39), with SICIEMG deficits observed in the former. Some individuals with ASD might also present atypical SICI_EMG_ responses, i.e. facilitation (40). In contrast, two studies applied LICI TMS-EMG protocols in ASD samples (39, 40), neither of which provide evidence for altered LICI_EMG_ response between ASD and neurotypical control groups. Again, however, both provide some evidence of individual variability in response to ppTMS protocols in the ASD group (39, 40). To the best of our knowledge, only two studies have reported TMS-EEG outcomes in this population to date, neither of which applied ppTMS or investigated GABA-ergic mechanisms (41, 42).

In the present study, single pulse (sp) TMS-EEG and ppTMS-EEG protocols were used to investigate the N100 component and overall LICI_EEG_ response of the TEP, respectively. Stimulation was applied to the right M1, right temporoparietal junction (TPJ), and right dorsolateral prefrontal cortex (DLPFC) in a group of adults with ASD (without intellectual disability) and matched neurotypical controls. The DLPFC and TPJ are widely implicated in the neuropathophysiology of ASD. Numerous studies demonstrate functional abnormalities at these regions (43–48), and they are also involved in neural networks implicated in the behavioural symptomatology of ASD, such as deficits in executive functions and theory of mind respectively (49–52). Given the wealth of knowledge that is available regarding TMS to M1, it was pertinent to include this site in the present investigation, along with TMS_EMG_ outcomes.

The two previous studies which investigated GABA_B_ mediated responses in ASD using TMS-EMG found no differences between ASD and neurotypical control participants (39, 40). EEG, however, provides a more direct measure of brain response to TMS, and as mentioned above, there is evidence against using ppTMS-EMG outcomes to infer ppTMS-EEG outcomes (33). Therefore, guided by our knowledge regarding the role of GABA-mediated cortical inhibition in ASD, we expected TMS-EEG indices of cortical inhibition (i.e., N100, LICI_EEG_) to be reduced in the ASD group compared to matched neurotypical controls, at all three target sites. LICI_EMG_ outcomes will also be reported.

## Methods

Data presented here were collected as part of an extensive multi-session study investigating the neuropathophysiology of ASD (15, 42, 53–55). Only information relevant to the reported data will be described herein. All protocols were approved by The Alfred Human Research Ethics Committee (213-11), and expedited approval was obtained from Monash University, Swinburne University, and Deakin University. All protocols were conducted in accordance with the Declaration of Helsinki. Participants were recruited via several means. Paper advertisements were placed around Monash and Swinburne University campuses, the Alfred hospital (Melbourne, VIC, Australia), and were also provided to local ASD community and support groups. Online advertisements were placed on social media platforms (e.g., Facebook and Gumtree). Participants were also recruited from the Monash Alfred Psychiatry Research Centre participant database. All participants provided written informed consent and were reimbursed a total of $50 AUD for their time.

### Participants

The sample presented here comprised 23 (11 males, 12 females) adults with ASD and 22 (11 males, 12 females) age, sex and IQ matched controls. Refer to Table 1 for descriptive statistics. According to the Edinburgh Handedness Inventory (56), all control participants were right-handed. In the ASD group, seven were left-handed, three ambidextrous, and 13 right-handed. Participants in the ASD group had been diagnosed by an external clinician (paediatrician, psychologist, or psychiatrist) before taking part in the study. A member of the research team saw a copy of each diagnostic report to confirm the diagnosis. All participants in the ASD group had been diagnosed based on DSM-IV criteria, which automatically assumes a DSM-5 diagnosis (1). To support diagnosis and characterise the behavioural phenotype of the sample, participants completed a battery of self-report measures related to traits and characteristics associated with ASD. Responses are summarised in Table 2.

**Table 1.** Participant demographics

|  | Control |  |  | ASD |  |  | U | *p* |
| --- | --- | --- | --- | --- | --- | --- | --- | --- |
|  | n | M (SD) | Median | n | M (SD) | Median |  |  |
| Age | 22 | 29.77 (9.79) | 27 | 23 | 30.30 (9.19) | 27.5 | 242 | 0.81 |
| Composite IQ<sup>1</sup> | 22 | 113.27 (14.27) | 116 | 22<sup>2</sup> | 110.41 (14.79) | 108.5 | 208 | 0.43 |

Note: <sup>1</sup> Composite IQ measured using the Kaufman Brief Intelligence Test, Second Edition (KBIT-2) (59). <sup>2</sup> Composite IQ score is not available for one participant in the ASD group due to fatigue related inattention.

**Table 2.**
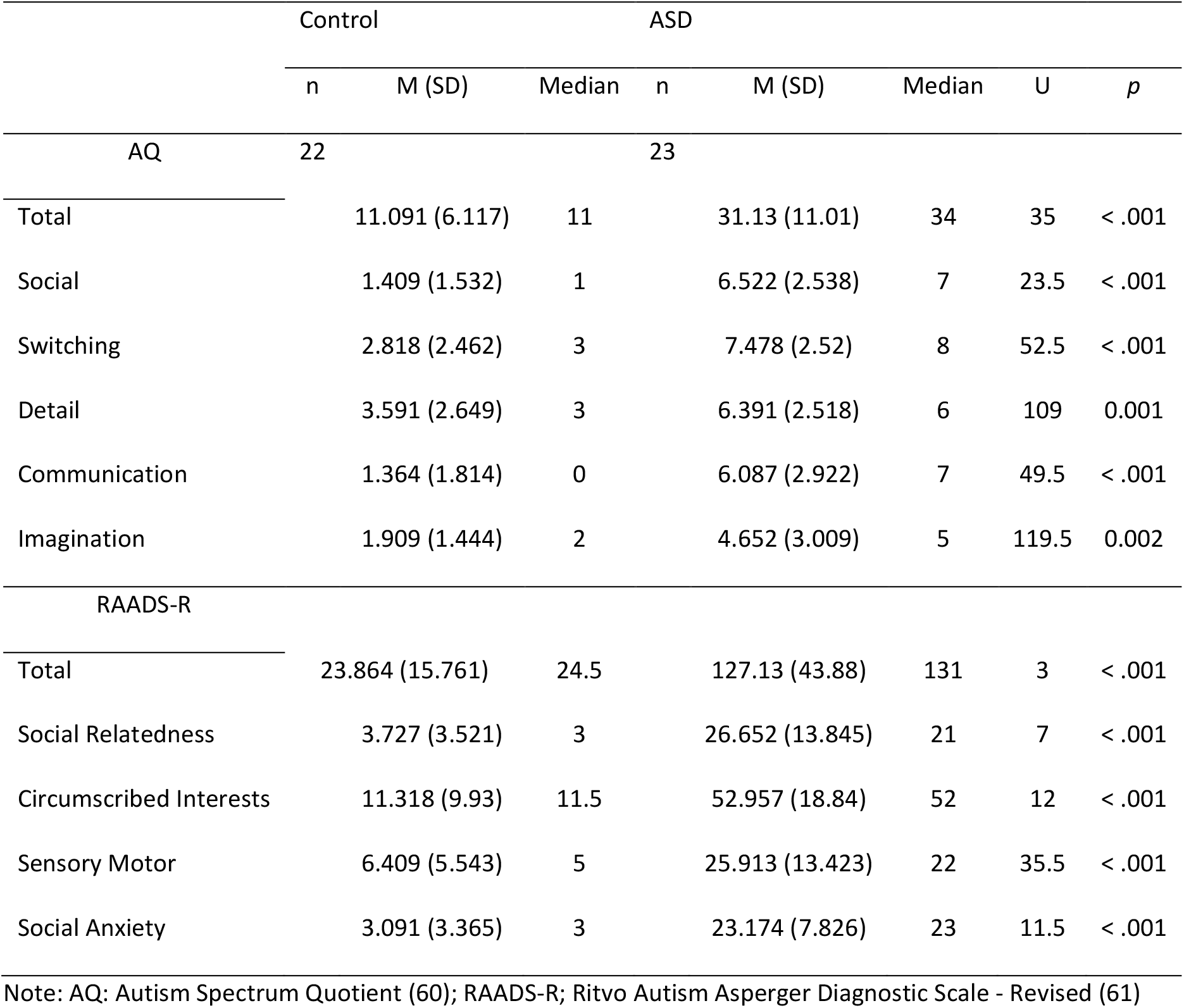
Summary of scores on self-report measures of traits and characterises associated with ASD

|  | Control |  |  | ASD |  |  | U | *p* |
| --- | --- | --- | --- | --- | --- | --- | --- | --- |
|  | n | M (SD) | Median | n | M (SD) | Median |  |  |
| AQ | 22 |  |  | 23 |  |  |  |  |
| Total |  | 11.091 (6.117) | 11 |  | 31.13 (11.01) | 34 | 35 | < .001 |
| Social |  | 1.409 (1.532) | 1 |  | 6.522 (2.538) | 7 | 23.5 | < .001 |
| Switching |  | 2.818 (2.462) | 3 |  | 7.478 (2.52) | 8 | 52.5 | < .001 |
| Detail |  | 3.591 (2.649) | 3 |  | 6.391 (2.518) | 6 | 109 | 0.001 |
| Communication |  | 1.364 (1.814) | 0 |  | 6.087 (2.922) | 7 | 49.5 | < .001 |
| Imagination |  | 1.909 (1.444) | 2 |  | 4.652 (3.009) | 5 | 119.5 | 0.002 |
| RAADS-R |  |  |  |  |  |  |  |  |
| Total |  | 23.864 (15.761) | 24.5 |  | 127.13 (43.88) | 131 | 3 | < .001 |
| Social Relatedness |  | 3.727 (3.521) | 3 |  | 26.652 (13.845) | 21 | 7 | < .001 |
| Circumscribed Interests |  | 11.318 (9.93) | 11.5 |  | 52.957 (18.84) | 52 | 12 | < .001 |
| Sensory Motor |  | 6.409 (5.543) | 5 |  | 25.913 (13.423) | 22 | 35.5 | < .001 |
| Social Anxiety |  | 3.091 (3.365) | 3 |  | 23.174 (7.826) | 23 | 11.5 | < .001 |

Note: AQ: Autism Spectrum Quotient (60); RAADS-R; Ritvo Autism Asperger Diagnostic Scale - Revised (61)

Participants were screened and excluded from this study based on standard contraindications to TMS (57). This includes seizure history, stroke, serious head injury, any brain-related illness or injury, ferromagnetic metal implants in the head, implanted medical devices such as pacemakers, pumps, or intracardiac lines, a history of persistent or severe headaches or migraines, medication which may increase risks associated with TMS, pregnancy, or lactation. Participants were also ineligible to partake in this study if they had a history of neurological or psychiatric illness (except for mood and anxiety disorders in the ASD group given the high prevalence among this population (58)). It is relatively commonplace for individuals with ASD to be prescribed psychotropic medications to manage difficulties associated with their diagnosis. Six participants in the ASD group reported taking one of the following psychotropic medications: serotonin-norepinephrine reuptake inhibitor (SNRI; n = 2), selective serotonin reuptake inhibitor (SSRI; n = 2), norepinephrine reuptake inhibitor (NRI; n = 1), and an atypical antipsychotic (n=1).

### Electroencephalography

Participants were fitted with a TMS compatible EEG cap (EASYCAP GmbH) and 20 silver-silver chloride (Ag-AgCl) sintered ring electrodes at the following 10-20 sites: AF4, F3, Fz, F2, F4, F6, FC4, C3, Cz, C2, C4, P3, Pz, P2, P4, P6, CP4, CP6, TP8, and PO4, as well as the bilateral mastoids; also refer to Figure 1. This montage was explicitly chosen as only electrodes surrounding our predetermined regions of interest (ROIs) for analysis, described in detail below, were deemed necessary. Electrooculogram (EOG) was recorded from electrodes placed at the outer canthus of each eye, and above and below the left eye. For EEG, electrodes were re-referenced online to CPz and the EOG electrodes to themselves. FCz served as the ground electrode. EEG recordings were conducted in a darkened room, amplified (x 1000), filtered (DC-2000 Hz), and digitised (10 kHz; SynAmps^2^, Compumedics Ltd.).

**Figure 1.**
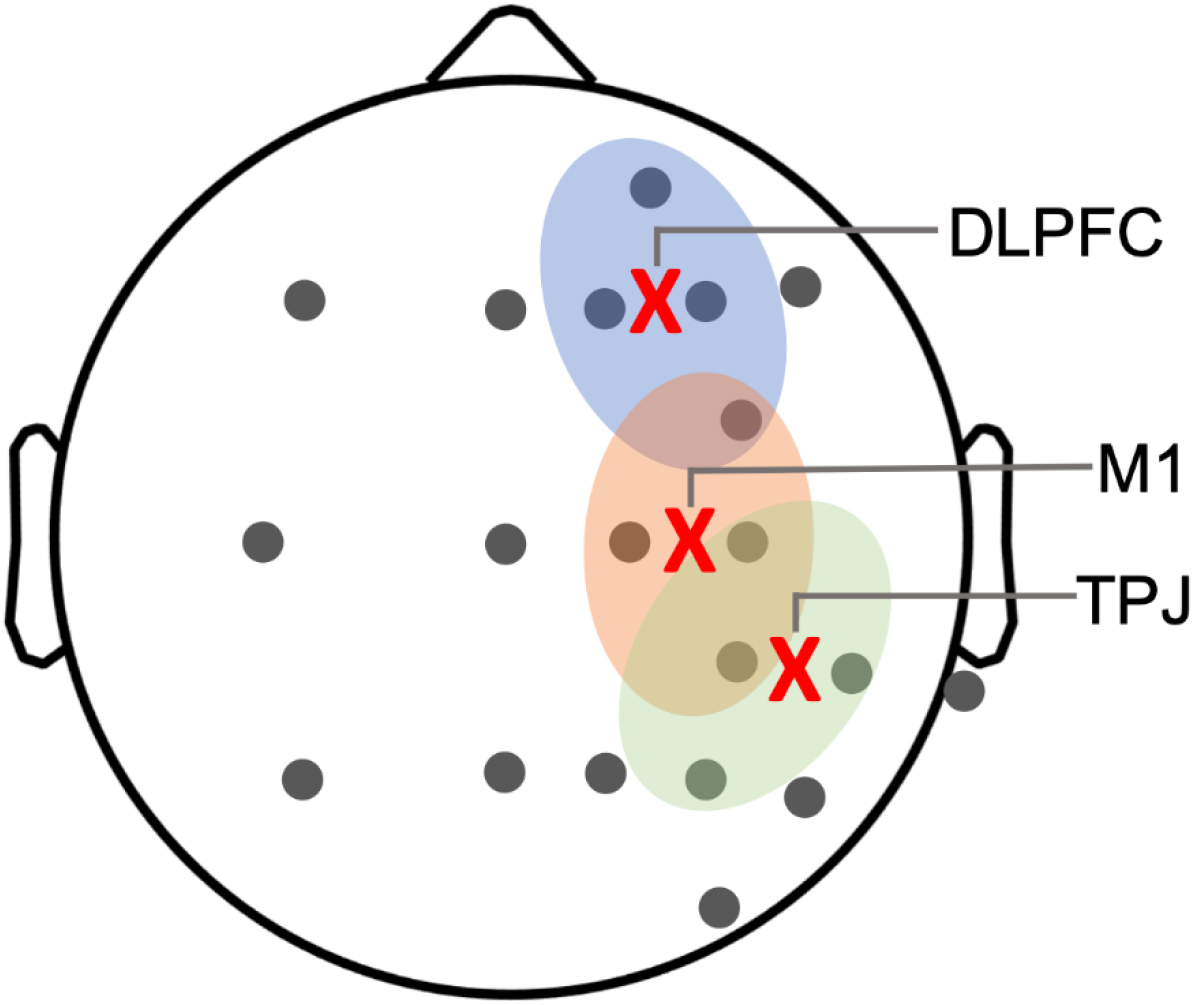
Graphical representation of electrode placement. The approximate TMS sites for each cortical target are marked with an ‘**X**’, EEG electrodes are in grey, and analysis regions of interest (ROIs) for each site are outlined (blue = dorsolateral prefrontal cortex [DLPFC], red = primary motor cortex [M1], green = temporoparietal junction [TPJ]).

### Transcranial Magnetic Stimulation

A figure-of-eight (70mm diameter) coil and two Magstim 200 stimulators, connected via a BiStim device (Magstim Ltd.), were used to deliver stimulation. All TMS was applied over the EEG cap.

### Motor threshold and electromyography

Participants were fitted with self-adhesive Ag-AgCl electrodes in a belly-tendon montage over the first dorsal interosseous (FDI) muscle, with a ground electrode placed over the ulnar styloid process, for EMG recording. The EMG signal, epoched around the TMS pulse (−200 to 500 ms), was amplified (x 1000) and band-pass filtered (10 - 500 Hz, PowerLab; ADInstruments) and digitised at 2 kHz with a Cambridge Electronic Design (CED Ltd.) interface.

Single TMS pulses, with the coil held at a 45° angle tangential to the midline, were delivered above the right M1 in a systematic pattern, with incremental increases in stimulator output, until the "hot spot", or site producing the strongest and most consistent MEPs at the FDI was identified. The stimulator output required to elicit an average MEP of 1 millivolt (mV) in peak-to-peak amplitude across ten consecutive trials (S1mV) was the stimulation threshold applied to each stimulation site for the experimental protocols. S1mV did not differ between groups; U = 182.5, *p* = .65. Due to recording error, S1mV is not available for 5 participants (3 control, 2 ASD).

### Experimental TMS protocol

Participants sat upright in a comfortable chair and were required to remain awake and keep their eyes opened, while blinking naturally, for the duration of the protocol. During stimulation, participants listened to white noise via disposable earphones inserted into the ear canal to attenuate auditory artefacts due to external noises, as well as the clicking sound elicited by the TMS pulses. All recording occurred in a darkened room to reduce interference from line noise. 75 single (see also: 42) and 75 paired TMS pulses were delivered randomly, 5 seconds apart with a 10% jitter. Stimulation was delivered consecutively to each site (M1, TPJ, DLPFC) in separate blocks, with the coil held 45° tangential to the midline at each site. Each block of stimulation lasted approximately 12 minutes, and participants were offered short breaks between blocks. For ppTMS, two S1mV TMS pulses were delivered 100 ms apart (i.e., conditioning pulse followed by a test pulse [LICI_100_]).

For M1 stimulation, TMS was delivered over the motor cortex hot spot. EMG was also recorded for this condition. For non-motor regions, TMS was delivered between CP4-CP6, and F2-F4 for the TPJ and DLPFC, respectively. These sites were chosen as CP6 (62) and F4 (63) have been used to guide TPJ and DLPFC stimulation previously in the absence of neuronavigation (62, 63). As the application of TMS laterally increases the likelihood of inducing muscle-related artefact (64), we placed the coil in a slightly more medial position. All EEG cables ran divergent to the coil, again to reduce TMS-related artefact (65).

### Data processing

#### EMG

ppTMS data (peak-to-peak amplitudes) were visually inspected for each trial in Signal 7.02, Cambridge Electronic Design, Cambridge, UK, and raw values were exported into Microsoft Excel (Office 365). Four EMG data files were corrupted and unable to be salvaged. Trials for which the conditioning pulse could not be visually detected, was below 0.4 mV, or was above 1.6mV were discarded. Trials for which there was evidence of tonic muscle activity prior to the TMS were also discarded. Data for three control participants were excluded due to tonic muscle activity being present throughout the entire block. The %change between the conditioning and test pulse in ppTMS trials was then calculated [(test pulse/conditioning pulse) x 100], and screened for individual outliers. Values exceeding three standard deviations away from the mean of %change data were removed. Only participants for whom 20 or more trials remained following this process were analysed (N = 34).

#### TMS-EEG Preprocessing

Both the spTMS and ppTMS data were analysed offline using Matlab (R2020a; The Mathwoks, MA, USA) incorporating the EEGLAB (66) and TESA (67) toolboxes. Participants data were excluded from a condition if there were <30 trials available for analysis. The final samples per block are clarified in Table 3. Data were segmented into 2000 ms epochs (−1000 to 1000 ms) around each TMS pulse (test pulse for the ppTMS data). The time window containing the high-amplitude magnetic TMS pulse artefact was removed and linearly interpolated (−5 to 15 ms), following which the data were corrected to the pre-stimulus baseline (−500 to −110 ms). Data were then down-sampled to 1 KHz, visually inspected, and any channels and/or trials containing excessive movement or muscle artefact were removed, refer to Supplementary Table S1. An initial round of ICA (FastICA; “tanh” contrast) (68) was then performed to remove large amplitude muscle artefacts caused by stimulation-induced contraction of the scalp muscles. Data were then band-pass (1-100 Hz) and bandstop (48-52 Hz) filtered (4^th^ order zero-phase Butterworth filter). The source-estimate-utilising noise-discarding (SOUND) algorithm (69) was then applied to further suppress TMS-evoked decay activity and any other noise-related activity (70). Data were again visually inspected, followed by a second round of ICA to remove any remaining artefacts (e.g., persistent muscle activity, eye blinks, lateral eye movements). Finally, any removed channels from the previous analysis steps were replaced using spherical interpolation and the data were re-referenced to the average of both mastoids. Both rounds of ICA incorporated semi-automatic component classification algorithms contained within the TESA software (TESA “compselect” function); all identified components were manually checked before being removed (67).

**Table 3.**
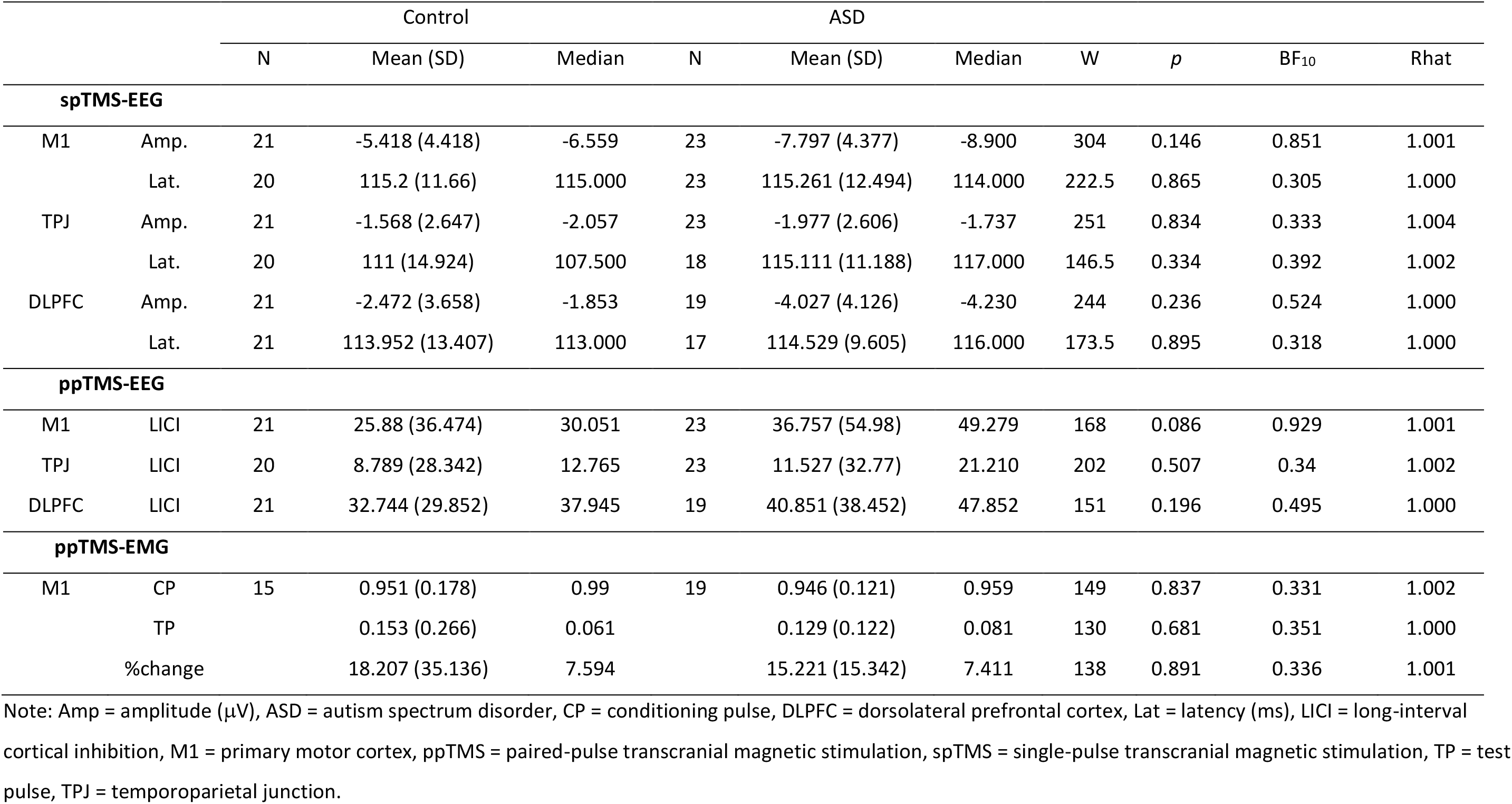
Summary of descriptive and inferential statistics for group comparisons

#### TMS-EEG Analysis

All TEP analyses were performed using data averaged across a group of four region of interest (ROI) electrodes close to the TMS site (Figure 1). For stimulation of M1, the ROI comprised the C2, C4, FC4, and CP4 electrodes; for DLPFC stimulation the ROI comprised the F2, F4, AF4, and FC4 electrodes; for TPJ stimulation the ROI comprised of CP4, CP6, C4, and P4 electrodes. For spTMS, N100 amplitudes across each of the M1, DLPFC, and TPJ ROIs for each subject and cohort (ASD and control) were extracted using the “*tesa_tepextract” and “tesa_peakanalysis”* functions within the TESA toolbox (67). The search window for detection of the N100 peak spanned 90-140 ms following the TMS pulse, i.e., the time window in which this peak typically occurs, and similar to previous studies (71–73). The N100 was defined as the largest negative deflection occurring within this window. In any instances where peaks were not detected within the specified window, amplitudes were extracted from the centre of the pre-defined latency windows. The average amplitude within ±5 ms either side of the detected peak was extracted and used for statistical analyses.

Consistent with previous studies, before calculating LICIEEG values, steps were taken to reduce contamination of the TEP produced by the test pulse in the ppTMS dataset by the conditioning pulse. Specifically, the spTMS data were time-shifted −100 ms to align with the conditioning pulse in the ppTMS dataset and were then subtracted from the ppTMS data to create an adjusted ppTMS response. This approach has been shown to significantly reduce the impact of the conditioning pulse on the TEP waveform (73–75). LICIEEG was then calculated using the formula:

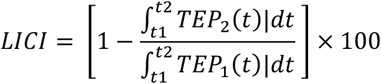

Where TEP_1_ refers to the evoked response from the spTMS, and TEP_2_ refers to the adjusted ppTMS response; while t1 and t2 refer respectively to the beginning and end of the time windows used for analysis. Consistent with past research, LICI_EEG_ was calculated for the entire TEP (50 – 275 ms) to explore cortical inhibition following the TMS pulse (75, 76).

## Results

### All statistical analyses were performed in JASP, Version 0.14

Raw data were found to violate the assumptions of normality. Several statistical outliers (i.e. values with z-scores exceeding 3) were identified in this data set, and these outliers were windsorised. This process did not improve the distribution of the data, and data transformations were then performed. Attempts at data transformation were not successful, and it was for this reason, as well as the relatively small sample size, that non-parametric Mann-Whitney U tests were performed. Missing data were excluded per dependant variable (pairwise), and all analyses performed were two-tailed.

There were no statistically significant differences between groups in the N100 amplitude or latency at any site following spTMS. Furthermore, LICI_EEG_ did not differ between groups based on TMS-EEG outcomes at any site. Regarding the TMS-EMG outcomes, again, our data provide no evidence of group differences in LICI_EMG_ at M1. Table 3 provides a summary of descriptive and inferential statistics. The TMS-EEG outcomes are also presented visually in Figures 2 (spTMS) and 3 (ppTMS), and the TMS-EMG data in Figure 4.

**Figure 2.**
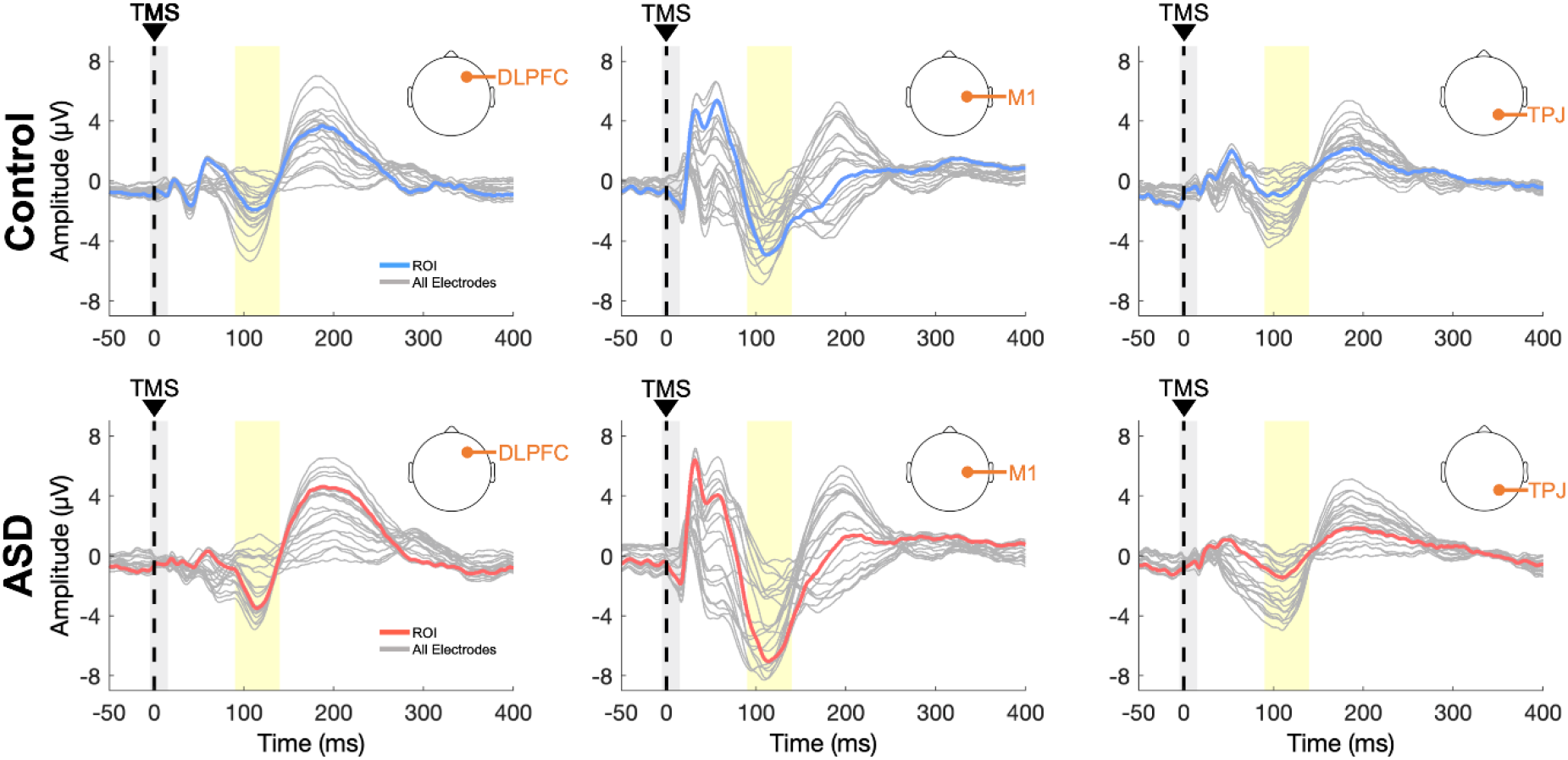
Butterfly plots demonstrating the TMS-evoked potential (TEP) waveform for the DLPFC, M1 and TPJ post single-pulse transcranial magnetic stimulation spTMS. The thick blue (control) and red (autism spectrum disorder; ASD) lines represent the TEP waveform as an average over the ROI electrodes for each stimulation site; the thin grey lines represent all other scalp electrodes. The vertical dashed line at 0 ms indicates the TMS pulse while the surrounding grey bar represents the −5 to 15 ms section of EEG containing the large TMS artefact which was removed and re-interpolated prior to analysis. The time-window used for detection of the N100 peak is denoted by the yellow shaded bar.

**Figure 3.**
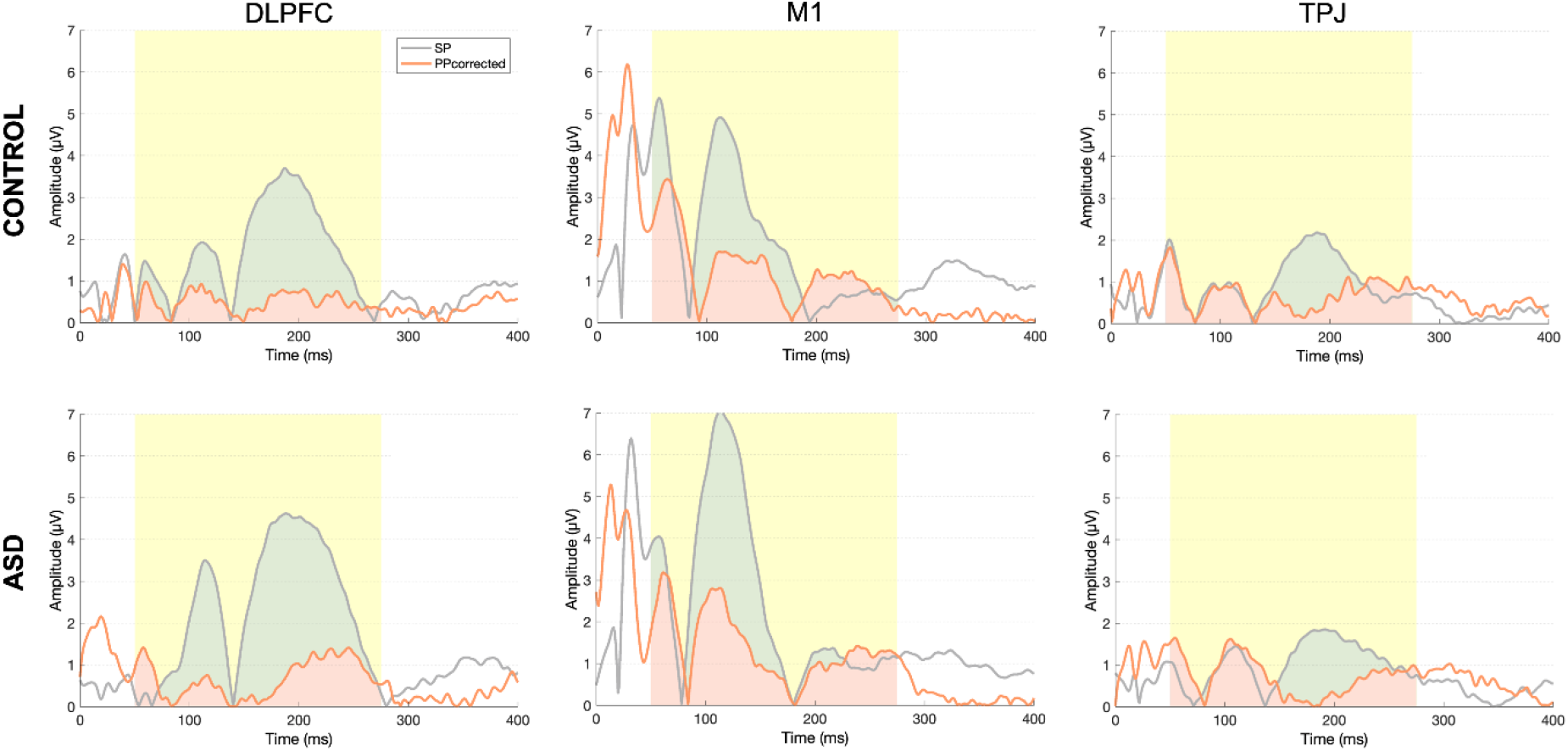
Graphical representation of long-interval cortical inhibition (LICI_EEG_). Rectified single-pulse (SP; grey) and corrected paired-pulse (PPcorrected; orange) waveforms following ppTMS-EEG are shown. The yellow shaded bar represents the time window used for calculation of LICI.

**Figure 4.**
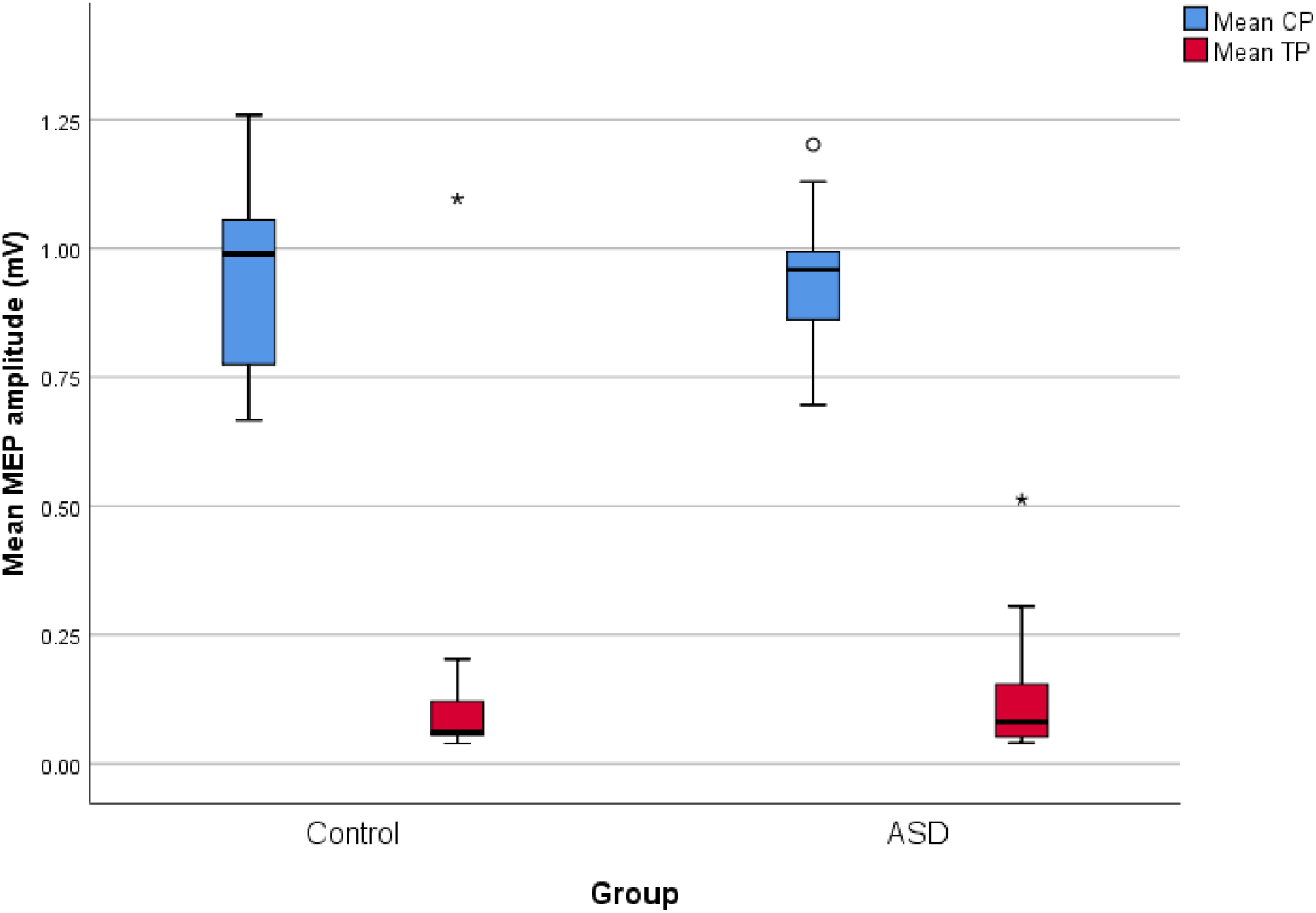
Box plots demonstrating the average MEP amplitude (mV) of the conditioning pulse (CP; blue) and test pulse (TP; red) for the control and ASD groups. * and ° represent statistical outliers in the TP and CP respectively, which were accounted for using non-parametric statistical analyses. A broader range of MEP amplitude is visible for the TP in the ASD group. To investigate this further, the coefficient of variation for the TP MEP was calculated (SD/M) for each participant and compared between groups. There was no difference in the variability of TP MEPs between groups, U = 108, p = .15, BF_10_ = 0.81.

To ensure that the findings identified by the frequentist analyses were not a consequence of data insensitivity, Bayesian Mann-Whitney U tests were then performed. This approach has been raised to be of particular importance to aid in elucidating whether null findings related to other NIBS (specifically theta-burst stimulation [TBS]) protocols are indeed due to a lack of effect, especially when dealing with smaller samples (77). The BF10 was used to test the likelihood of the data supporting either the null or alternative hypotheses. A BF_10_ value <1 would indicate that the data are in favour of the null hypothesis, while a BF_10_ value >1 would indicate that the data are in favour of the alternative hypothesis. Specifically, a BF_10_ between 0.33-1 is considered to provide weak support for the null hypothesis, BF_10_ between 0.33-0.03 provides moderate support for the null hypothesis, and a BF_10_ <0.03 provides strong support for the null hypothesis. Conversely, a BF10 value between 1-3 provides weak support for the alternative hypothesis, a BF10 value between 3-10 provides moderate support for the alternative hypothesis, and a BF_10_ value >10 provides strong support in favour of the alternative hypothesis (78, 79). For all comparisons, our models provide weak-moderate support in favour of the null hypothesis, and therefore support the outcomes of our frequentist analyses. That is, neither N100 amplitude nor latency following spTMS, or ppTMS induced LICI, as measured by TMS-EEG or TMS-EEG, differed between the ASD and control groups. The results of these analyses are also provided in Table 3.

## Discussion

We present the first study, to our knowledge, to implement a TMS-EEG paradigm to investigate cortical inhibition in ASD. spTMS and ppTMS were applied to the right M1, right TPJ, and right DLPFC in a sample of participants with ASD (without intellectual disability), and a matched control group. It was hypothesised that the N100, a TEP peak indirectly reflective of GABA_B_-ergic processes would be reduced in the ASD group, indicative of less cortical inhibition. The results of this present study did not support this hypothesis. It was further predicted that LICI_EEG_ would demonstrate less inhibition of the TEP among the ASD group. Again, this hypothesis was not supported. In line with the previous, albeit limited, literature using LICI ppTMS-EMG protocols to investigate cortico-spinal inhibition in ASD, ppTMS to M1 did not affect LICI_EMG_ outcomes.

The literature investigating the role of GABA in the neuropathophysiology of ASD is highly variable. While there is considerable MRS evidence to suggesting that GABA concentration is reduced in ASD (80, 81), a growing body of literature using TMS to investigate GABA-related synaptic activity challenges this (3, 37). It is important to emphasise, however, that MRS and TMS index different GABA-ergic properties (i.e., metabolite concentration and synaptic activity, respectively), and there is no evidence of a relationship between MRS- and TMS-related indices of GABA-ergic mechanisms, particularly concerning LICI_EMG_ (82, 83). Nevertheless, the data presented here overlap with a previously published study from our group, which reports on GABA-related MRS outcomes (15). A sub-set of participants from this study underwent MRS, acquired from voxels placed at the right DLPFC and right TPJ, and GABA concentration was found not to differ between groups. We must, therefore, not discount the possibility that the observed results of this study are in some way specific to the sample. The extent to which they might apply to other ASD samples, including younger individuals, is unclear. Indeed, there is some TMS-based evidence to suggest that GABA-ergic impairments might be more readily observable in individuals with ASD presenting with more severe impairment (e.g., language; (38)).

Regarding TMS, there is also a growing body of evidence demonstrating vast individual variability regarding various TMS-related outcomes (84–86). Indeed, Oberman and colleagues (40) report facilitatory effects of both SICI and LICI protocols in some individuals with ASD. While the mechanisms underlying this variability remain largely unknown, several factors such as age, [epi]genetics, and biological sex are likely contributors (87, 88), as is the target/stimulation location and tasks/state at the time of outcome recording (89). When considering the application of TMS in the field of ASD, these factors are of particular importance as age (90), biological sex (91, 92), and genetics (93, 94) are all known to contribute to the vast heterogeneity of this condition. Notably, some of the genes known to affect the response to non-invasive brain stimulation, including TMS (95), are also implicated in ASD (96, 97). More work in this domain is imperative.

## Limitations

There are some limitations of this study that must be acknowledged. Firstly, our sample is relatively small; however, the use of non-parametric and Bayesian analyses assist in mitigating this factor. More importantly, regarding the sample, here we present data only for adults without intellectual disability. An essential consideration in this regard is that age-related decline in GABA concentrations has been reported in healthy adults (98), and it is unclear whether this same effect occurs in ASD. These data, therefore, cannot be generalised to younger samples, or those with more severe impairment. An important methodological limitation is that stereotaxic neuronavigation was not available for this study. The target sites are, however, based on previous research also targeting these regions. Finally, while we did use white-noise to mitigate the impacts of auditory artefacts, somatosensory (tactile) effects were not controlled for in the present study. Recent research indicates that somatosensory artefacts might have effects across the TEP (99), and can specifically impact later TEP peaks (100, 101), including the N100. This, therefore, must be considered when interpreting these results.

## Conclusions

To conclude, this study applied spTMS-EEG and ppTMS-EEG to the right M1, right TPJ, and right DLPFC in a sample of adults with ASD and matched neurotypical controls. Using this method, the results of this study do not provide evidence to indicate GABAB-ergic deficits in this sample. Care must be taken when interpreting these findings as generalisability is limited. Further research investigating these mechanisms at various ages and developmental stages, as well as in individuals with various levels of symptom severity is needed. These protocols could also be incorporated with pharmaceutical trials investigating the therapeutic potential of GABA-ergic agonists in ASD to understand further the effects of such drugs at cortical regions implicated in ASD. Factors contributing to variability in TMS outcomes, in clinical and non-clinical populations, must also be elucidated.

## List of Abbreviations

ASD: autism spectrum disorder
DLPFC: dorsolateral prefrontal cortex
EEG: electroencephalography
EMG: electromyography
EOG: electrooculogram
FDI: first dorsal interosseous
GABA: gamma-aminobutyric acid
ISI: inter-stimulus interval
LICI: long interval cortical inhibition
M1: primary motor cortex
MRS: magnetic resonance spectroscopy
NIBS: non-invasive brain stimulation
NRI: norepinephrine reuptake inhibitor
ppTMS: paired-pulse transcranial magnetic stimulation
ROI: region of interest
SICI: short interval cortical inhibition
SNRI: serotonin-norepinephrine reuptake inhibitor
spTMS: single-pulse transcranial magnetic stimulation
SSRI: selective serotonin reuptake inhibitor
TEP: transcranial magnetic stimulation-evoked potential
TMS: transcranial magnetic stimulation
TPJ: temporoparietal junction

## Declarations

### Ethics approval and consent to participate

All protocols were approved by The Alfred Human Research Ethics Committee (213-11), and expedited approval was obtained from Monash University, Swinburne University, and Deakin University. All protocols were conducted in accordance with the Declaration of Helsinki. All participants provided written informed consent.

### Consent for publication

Not applicable

### Availability of data and material

Deidentified data may be accessed from the corresponding author upon reasonable request, and following approval from relevant Human Research Ethics Committees/Institutional Review Boards.

### Competing interests

PBF has received equipment for research from MagVenture A/S, Medtronic Ltd, Neuronetics and Brainsway Ltd and funding for research from Neuronetics. He is on scientific advisory boards for Bionomics Ltd and LivaNova and is a founder of TMS Clinics Australia. The authors report no other conflicts of interest.

### Funding

MK and AH are supported by Alfred Deakin Postdoctoral Research Fellowships. BMF was supported by a NHMRC Early Career Fellowship (1070073). PBF is supported by a NHMRC Practitioner Fellowship (1078567). PGE is supported by a Future Fellowship from the Australian Research Council (FT160100077).

### Authors Contributions

MK and PGE contributed to conception of the work, study design, data acquisition and analysis, interpretation of results, and preparation of manuscript. AH contributed to the data analysis, interpretation of results, and preparation of manuscript. NCR contributed to the study design, data acquisition and analysis. TS and BMF contributed to data acquisition. JY, MD, PHD and NAU contributed to data analysis and interpretation of results. PBF contributed to the conception of the work and study design. All authors have read, contributed to, and approve the final manuscript.

## Acknowledgements

We thank all the participants who volunteered their time to take part in this research.

