## Supplementary Table S1 for "A single- and paired-pulse TMS-EEG investigation of the N100 and long interval cortical inhibition in autism spectrum disorder"

Table S1. Average remaining TMS-EEG trials and channels after preprocessing (± SD)

| | DLPFC | | M1 | | TPJ | |
|---|---|---|---|---|---|---|
| | spTMS | ppTMS | spTMS | ppTMS | spTMS | ppTMS |
| Controls | | | | | | |
| Remaining Trials | 69.33 (11.32) | 67.81 (12.79) | 74.52 (2.83) | 70.95 (6.57) | 73.95 (2.95) | 74.15 (1.92) |
| Remaining Channels | 18.00 (0.00) | 17.95 (0.22) | 17.90 (0.30) | 17.85 (0.36) | 17.90 (0.30) | 17.95 (0.22) |
| ASD | | | | | | |
| Remaining Trials | 74.47 (2.76) | 72.74 (3.69) | 74.34 (3.89) | 73.13 (5.14) | 70.96 (4.43) | 72.74 (4.97) |
| Remaining Channels | 17.95 (0.23) | 17.89 (0.32) | 17.83 (0.39) | 17.91 (0.29) | 17.78 (0.52) | 18.00 (0.00) |
